# Zika virus-based immunotherapy enhances long-term survival of rodents with brain tumors through upregulation of memory T-cells

**DOI:** 10.1101/2020.04.24.059253

**Authors:** Andrew T. Crane, Matthew R. Chrostek, Venkatramana D. Krishna, Maple Shiao, Nikolas G. Toman, Clairice Pearce, Sarah K. Tran, Christopher J. Sipe, Winston Guo, Joseph P. Voth, Shivanshi Vaid, Hui Xie, Wei-Cheng Lu, Will Swanson, Andrew W. Grande, Mark R. Schleiss, Craig J. Bierle, Maxim C-J. Cheeran, Walter C. Low

## Abstract

Zika virus (ZIKV) exhibits a tropism for brain tumor cells and has been used as an oncolytic virus to target brain tumors in mice with modest effects on extending median survival. Recent studies have highlighted the potential for combining virotherapy and immunotherapy to target cancer. We postulated that ZIKV could be used as an adjuvant to enhance the long-term survival of mice with malignant glioblastoma and generate memory T-cells capable of providing long-term immunity against cancer remission. To test this hypothesis mice bearing malignant intracranial GL261 tumors were subcutaneously vaccinated with irradiated GL261 cells previously infected with the ZIKV. Mice also received intracranial injections of live ZIKV, irradiated ZIKV, or irradiated GL261 cells previously infected with ZIKV. Long-term survivors were rechallenged with a second intracranial tumor to examine their immune response and look for the establishment of protective memory T-cells. Mice with subcutaneous vaccination plus intracranial irradiated ZIKV or intracranial irradiated GL261 cells previously infected with ZIKV exhibited the greatest extensions to overall survival. Flow cytometry analysis of immune cells within the brains of long-term surviving mice after tumor rechallenge revealed an upregulation in the levels of T-cells, including CD4^+^ and tissue-resident memory CD4^+^ T-cells, in comparison to long-term survivors that were mock-rechallenged, and in comparison to naïve untreated mice challenged with intracranial gliomas. These results suggest that ZIKV can serve as an adjuvant to subcutaneous tumor vaccines that enhance long-term survival and generate protective tissue-resident memory CD4^+^ T-cells.

## Introduction

Glioblastoma multiforme (GBM) is a highly aggressive malignant brain tumor whose treatment options currently offer little chance of long-term survival or cure. Patients with GBM typically undergo surgical resection followed by treatment with chemotherapy and temozolomide (1), with a median overall survival (OS) of only 16 months (2). Given the limitations of available therapies, new approaches to treat GBM are needed.

Oncolytic viruses offer a promising avenue for treating GBM. Viral tropism can be exploited to target tumor cells with greater efficiency and fewer side effects when compared to treatment with chemotherapy and temozolomide (3). Recently, virotherapy has turned towards the immunogenic activity of viruses and their potential for immunotherapy (4). Viruses are thought to induce an anti-tumor immune response through the direct lysis of infected cells and release of tumor-associated antigens, as well as the activation of antiviral pathways, creating an adaptive immune response against tumor and virus (5).

In the present study, we investigated Zika virus (ZIKV) as a virotherapy for GBM. ZIKV has been shown to exhibit a tropism for GBM (6,7) and in mouse GBM models treatment with different viral strains, including a mouse-adapted Dakar ZIKV and a genetically modified Cambodian ZIKV, have improved OS (6,8). Here the therapeutic potential of the French-Polynesian (H/PF/2013) strain of ZIKV was assessed in a mouse model of glioma using the GL261 cell line. While ZIKV was capable of infecting GL261 cells *in vitro*, we observed no significant improvement in OS following intratumoral injections of ZIKV in GL261-bearing mice, unlike the survival improvements in a previous publication which used a mouse-adapted Dakar ZIKV to treat GL261-bearing mice (6). An increase in pattern recognition receptor transcripts was observed following *in vitro* ZIKV infection of GL261 cells, however, which led us to interrogate ZIKV as an adjuvant to vaccine-based immunotherapy. It was observed that intratumoral treatment with live or gamma-irradiated (IR) ZIKV in combination with repeated vaccination of IR tumor cells previously infected with ZIKV significantly improves OS in GL261-mice. Additionally, we provide evidence of enhanced T-cell response in the brain of mice surviving long-term after tumor induction, specifically CD4^+^ and memory CD4^+^ T-cells, suggestive of long-term immunity against glioma.

## Methods

All methods involving the use of ZIKV and mice described here have been approved by the University of Minnesota Institutional Biosafety Committee (Protocol 1910-37492H) and the Institutional Animal Care and Use Committee (Protocol 1910-37491A). Any work involving ZIKV was performed under BSL2 containment.

### Cell Culture

Mouse glioma cell line GL261-GFP.Luciferase (GL261; acquired from the lab of Dr. John Ohlfest), rat glioma cell line GS-9L (9L; ECACC, 94110705), mouse microglia BV2 cell line (acquired from the lab of Dr. Ling Li), and Vero cell line (African Green Monkey kidney epithelium; ATCC, CCL-81) were maintained with media changes every other day and cells were passaged when reaching 80% confluence using TrypLE. Glioma media consisted of DMEM high glucose and L-glutamine (Genesee Scientific 25-500), supplemented with 10% Fetal Bovine Serum (Corning 35-011-CV), 1% Penicillin-Streptomycin (HyClone SV30010), 1% MEM NEAA (Gibco 1140-050). Vero media consisted of MEM Earle Salts supplemented with L-glutamine (Genesee Scientific 25-504), 10% Fetal Bovine Serum (Corning 35-011-CV), 1% Penicillin-Streptomycin (HyClone SV30010), 1% MEM NEAA (Gibco 1140-050).

### ZIKV

ZIKV H/PF/2013 (passage 4) was obtained from the European Virus Archive and cultured using previously established protocols (9). ZIKV was passaged twice on Vero cells to generate working stocks of virus which were then concentrated by the ultracentrifugation of virus-containing media over a sucrose cushion, as described previously (10). ZIKV titers were calculated by titration and plaque assay. For infection, ZIKV was diluted in PBS to achieve the desired infectious dose. To make irradiation-inactivated ZIKV (irZIKV), the desired concentration of ZIKV was irradiated at 60 Gy with gamma irradiation from a Cs-137 irradiator.

### *In vitro* ZIKV infections

Briefly, 2-3 × 10^6^ cells (GL261) were infected with ZIKV at an MOI of 0.01 for two hours. Following incubation, the virus containing media was removed, the cells were rinsed with PBS to remove virus not adsorbed, and 4mL of fresh non-virus containing media was placed on the cells. Following the infection, virus-containing media was collected at 1-, 2-, and 3-days post-infection (DPI) and stored at −80°C. ZIKV concentrations from this media were calculated by plaque assay (11).

Plaque assays were performed to determine the amount of virus in the supernatant at each time point. A single six-well plate of Vero cells was made for each aliquot of supernatant. Approximately 300,000 Vero cells were plated in each well the day before infection and allowed to form a monolayer. The day after creating plates, viral supernatant was thawed and 10-fold serial dilutions were prepared from 10-1 to 10-6. 1 mL of each dilution was placed on each well and allowed to adsorb for 2 hours. After the adsorption period, a PBS wash was conducted to remove remaining virus not adsorbed, and finally a solution of 1.5mL of 2x concentrated Vero media and 1.5mL of 1.1% SeaPlaque low-melting agarose at 37oC was applied over the monolayers. This mixture was allowed to cool to room temperature, forming a gelatinous overlay. After four days, 4% PFA was applied for a minimum of 2 hours to fix the virus, cells, and overlays. The overlays were removed by applying warm tap water and manually tapping the plate, and 0.1% crystal violet was used to stain the cells and easily identify plaques. Plaques were counted under a dissection microscope and the concentration of virus in each day’s supernatant was calculated. Averages of each respective triplicate were used to calculate the concentration of virus at each time point for each cell line.

### RNA isolation

For RNA isolation, 2-3 × 10^6^ cells (GL261 and BV2) were infected with ZIKV at an MOI of 0.01 for two hours. Virus containing media was removed, cells were washed with PBS to remove any residual virus, and fresh media was added. Every 24 hours following infection, for three days, cells were lysed in RLT buffer and stored at −80°C before RNA isolation. Control samples of uninfected GL261 and BV2 cells were also collected. RNA was isolated from infected (1-DPI and 3-DPI) and uninfected GL261 and BV2 cells using the Qiagen RNeasy Plus Mini Kit following manufacturers’ instructions. Cell counts were performed in ZIKV-infected and uninfected cells and RNA was normalized to cell counts post-extraction.

### cDNA synthesis and qRT-PCR

cDNA was synthesized from 500 ng of each respective RNA sample using the ProtoScript® First Strand cDNA Synthesis Kit (New England BioLabs, E6300L) according to manufacturer’s instructions.

Using the Eppendorf Realplex 2 PCR system, qRT-PCR was performed using 100 ng of cDNA with 2.5 µl of specific primers, targeting MDA5, RIG-I, STAT2, IFN-β, TLR3 (Supplemental Table S1), and 12.5 µl of Apex qPCR 2X GREEN Master Mix (Apex Bioresearch Products) in a final reaction volume of 25 µl. The cycling conditions for MDA5, RIG-I, and STAT2 were 40 cycles of 95 °C for 30 s, 55 °C for 40 s, and 72 °C for 30 s. The cycling conditions for IFN-β and TLR3 were 40 cycles of 95 °C for 30 s, 50 °C for 40 s, and 72 °C for 30 s. mRNA expression fold change (over uninfected cells) was quantified by calculating 2-ΔΔCT with HPRT mRNA as an endogenous control.

### Tumor vaccine

Vaccines consisted of IR GL261 cells previously infected with ZIKV, coupled with GM-CSF. Briefly, cells were incubated with 0.5 MOI ZIKV for 48 hours, after which the media was removed and replaced with fresh media. The cells were cultured for an additional 48 hours followed by dissociating the culture monolayer with TrypLE and to obtain a cell pellet. The pellet was resuspended in PBS and, using a Cs-137 irradiator, irradiated with gamma rays for 60 Gy. The irradiated cell solution was centrifuged, then resuspended in PBS and total cell number was counted using a hemocytometer. The cells were centrifuged and resuspended in freezing media (glioma media with 10% DMSO) at a concentration of 1 × 10^7^ cells/mL/cryovial. The cryovial was frozen in isopropanol chambers at −80 oC until use. On days 3, 7, and 14 or 21 (for the first and second treatment studies, respectively) following tumor-induction, cryovials were removed from −80 oC, thawed in a 37 oC water bath, then resuspended in warm glioma media. Cells were centrifuged, washed in glioma media followed by another centrifugation. The resulting cell pellet was then resuspended in 1mL PBS with 20ng GM-CSF. Mice in the vaccination groups received a subcutaneous injection of 500 µL.

### Preparation of GL261 for tumor induction

Prior to surgery, GL261 cells were prepared for transplantation by washing three times with PBS, then trypsinized for 5 minutes at 37°C. The cell suspension was collected then centrifuged and the resulting pellet was resuspended in cold DMEM for counting viable cells using the Trypan Blue exclusion. The cells were centrifuged a second time and resuspended at a concentration of 1 × 10^4^ cells per µL in cold DMEM and placed on ice.

### Mouse tumor induction and treatment

10-week old C57BL/6J and NOD-SCID mice purchased from Jackson Laboratories were housed in environmentally controlled micro-isolator cages with Enviro-Dry environmental enrichment. Mice were first anesthetized with isoflurane oxygen mixture, then the head of the animal was shaved and treated with betadine. Following mounting in a stereotaxic frame, a single midline incision was made along the scalp and skin retracted to expose bregma. A small burr hole was drilled in the skull above the injection site in the right hemisphere (from bregma: anterior 1.0 mm and lateral 1.5 mm). A 10 µL Hamilton syringe loaded with the cell solution was slowly inserted into the brain (3.1 mm ventral to the pia mater in mice) and 1 × 10^4^ viable cells in 1 µL were injected at a rate of 0.5 µL/minute. Following injection, the needle remained in place for one minute. The injection needle was raised 0.1 mm then 0.2 mm from the initial injection site and the injection was repeated with 1 × 10^4^ cells injected at each site for a total of 3 × 10^4^ viable cells (total volume of 3 µL). At the conclusion of the last injection, the needle remained in place for three minutes before being slowly withdrawn.

Mice scheduled for treatment with intracranial (*i.c.*) ZIKV or irZIKV immediately following tumor induction remained in the stereotaxic frame. A 10 µL Hamilton syringe was loaded with 5 × 10^4^, 5 × 10^6^, or 5 × 10^8^ pfu/µL of ZIKV or 5 × 10^4^ irZIKV. The needle was slowly inserted into the brain (3.0 mm) and 1 µL of virus was injected at a rate of 1 µL/minute. At the conclusion of the injection, the needle remained in place for one minute before being slowly withdrawn. Mice not treated with *i.c.* ZIKV were injected with sterile saline at the same coordinates. At the conclusion of the surgery, the incision site was cleaned and closed using a 7mm Reflex wound stapler. Mice were transferred to a heated recovery cage until fully sternal, then returned to their homecage.

Mice scheduled for treatment with *i.c.* irZIKV or *i.c.* vaccine 14-days post-tumor induction were anesthetized and prepped for surgery as described above with the center of the burr hole from the prior surgery serving as the injection site. The needle was slowly inserted into the brain (3.25 mm ventral to the pia mater in mice) and 1 µL, containing 5 × 10^4^ pfu/µL irZIKV or 3 × 10^4^ IR GL261 cells previously infected with ZIKV, was injected at a speed of 0.5 µL/minute.

Following injection, the needle remained in place for one minute. The injection needle was raised 0.75 mm from the initial injection site and the injection was repeated, totaling 1 × 10^5^ pfu/µL irZIKV or 6 × 10^4^ IR GL261 cells previously infected with ZIKV. At the conclusion of the last injection, the needle remained in place for three minutes before being slowly withdrawn. Mice were transferred to a heated recovery cage until fully sternal, then returned to their homecage.

Following tumor induction, mice were monitored by lab member and University of Minnesota veterinary technicians three times daily, for up to 118-days post-tumor induction. Moistened lab chow was placed in a dish in the bedding to allow mice access to food and fluids. The point at which the mice were unable regain an upright posture, or demonstrate response to physical stimulation from the observer, or unable to lift their neck and head, the animal was declared moribund and immediately euthanized via Ketamine overdose (100mg/kg, *i.p.*) followed transcardial perfusion with ice-cold PBS and 4% paraformaldehyde.

### Tumor re-challenge

Long-term survivors (118-days and 86-days following initial tumor induction for the first and second study, respectively) were injected with 3 × 10^4^ GL261 cells in both the left- and right-hemispheres, following the procedure described above. Seven days following rechallenge, mice were deeply anesthetized with an overdose of Ketamine (100mg/kg, *i.p.*) and. The brain was removed and the right hemisphere was extracted and dissociated for flow cytometry. Control mice included long-term survivors with an intracerebral injection of saline, age-matched controls with *i.c.* tumor, and age-matched controls with *i.c.* saline injection.

### Flow cytometry

Mononuclear cells isolated from tumor re-challenged mouse brains were isolated by a density gradient (12). Briefly, brains were finely minced using razor blades, mechanically disrupted by pipetting up and down, and suspended in RPMI medium with 30% Percoll. This suspension was transferred to 15 ml conical tube and slowly underlayed with 1.5 ml of 70% Percoll solution. After centrifugation at 900 x g for 30 min at 15 oC, the mononuclear cells in the interphase were collected and washed twice with PBS. Cells were stained with LIVE/DEADTM fixable Aqua dead cell stain (Thermo Fisher Scientific, Rockford, IL) for 30 min at 4oC, blocked with anti-mouse CD16/CD32 (Fc block, BD Biosciences) for 5 min at RT, and stained with following anti-mouse immune cell markers for 30 min at 4 oC. CD45-PerCP-Cyanine5.5, CD11b-APC-eFluor 780, Ly6c-eFluor 450, CD86-FITC, MHC class II-PE, CD19-PE, CD3e-eFluor 450, CD4-PE-Cy7, CD8-FITC, CD44-APC, CD25-PE, (eBioscience, San Diego, CA), CD11c-Brilliant Violet 711, Ly6G-PE-Cy7, CD206-Alexa Fluor 647, CD62L-Brilliant violet 605, NK1.1-Brilliant violet 650, CD69-Brilliant violet 711, (BioLegend, San Diego, CA). To identify Tregs, surface stained cells were fixed, permeabilized and stained with FoxP3-APC using anti-mouse/rat FoxP3 staining set (eBioscience, San Diego, CA). Isotype specific antibodies were used to control for nonspecific antibody binding. Immunostained cells were acquired using BD LSRFortessa X-20 flow cytometer (BD Biosciences) and data were analyzed using FlowJo software. SPHERO AccuCount particles (Spherotech, Lake Forest, IL) were added to samples immediately before analysis to calculate absolute number of each cell population.

### *In vitro* stimulation and IFN-γ detection

Cervical lymph nodes (CLN), from rechallenged long-term surviving mice treated with *i.c.* vaccines or control mice were mechanically disrupted to prepare single-cell suspension and passed through 70 µm cell strainer. Cells were washed twice with PBS and resuspended in complete RPMI medium (RPMI 1640 medium supplemented with 10% FBS, 2mM L-glutamine, 1mM sodium pyruvate, 20mM HEPES, and 1x Penicillin-Streptomycin). For IFN-γ assay, 0.5×10^6^ live cells were seeded in 200 µL per well in a 96-well flat bottom tissue culture plate and stimulated in triplicate wells with irradiated GL261 cells lysate (50µg/mL) or heat-inactivated ZIKV (3.5×10^6^ pfu/ml). Cells stimulated with Concanavalin A (10 µg/ml) or unstimulated cells were included as positive and negative controls respectively. Cells were incubated at 37oC/5% CO_2_ for 72 h. Culture supernatants were collected at 72 h of stimulation and concentration of interferon-γ (IFN-γ) was determined using mouse IFN-gamma DuoSet ELISA (DY485, R & D systems, Minneapolis, MN) according to manufacturer’s instructions.

### Rat tumor induction and treatment

11-week old F344 rats weighing 225-250g purchased from Taconic were housed in environmentally controlled micro-isolator cages with Enviro-Dry environmental enrichment. Animals were first anesthetized with isoflurane oxygen mixture, then the head of the animal was shaved and treated with betadine. Following mounting in a stereotaxic frame, a single midline incision was made along the scalp and skin retracted to expose bregma. A 10 µL Hamilton syringe was loaded with the 9L cell solution (prepared for tumor induction following the mouse protocol above). A small burr hole was drilled in the skull above the injection site in the right hemisphere (from bregma: anterior 1.0 mm and lateral 3.0mm). The needle was slowly inserted into the brain (5 mm ventral to pia mater in rats) and 2.5 × 10^4^ viable cells were injected at a speed of 0.5 µL/minute. Following injection, the needle remained in place for one minute. The injection needle was raised 0.1 mm from the initial injection site and the injection was repeated with 2.5 × 10^4^ cells injected at each site for a total of 5 × 10^4^ viable cells. At the conclusion of the last injection the needle remained in place for three minutes before being slowly withdrawn. Animals scheduled for treatment with intra-tumoral (IT) ZIKV were transferred to a BSL-2 laminar flow hood while remaining under isoflurane anesthesia in the stereotaxic frame. A 10 µL Hamilton syringe was loaded with ZIKV or IR ZIKV at viral titers of 5 × 10^4^ pfu/µL. The needle was slowly inserted into the brain (4.5 mm) and 1 µL of virus was injected at a speed of 1 µL/minute. At the conclusion of the injection the needle remained in place for one minute before being slowly withdrawn. Animals not treated with ZIKV were injected with sterile saline. At the conclusion of the surgery, the incision site was cleaned and closed using a 9mm Reflex wound stapler. Animals were transferred to a heated recovery cage until fully sternal. Vaccines consisted of IR 9L cells previously infected or uninfected with ZIKV, coupled with GM-CSF. Briefly, cells were incubated with 0.5 MOI ZIKV for 48 hours, after which the media was removed and replaced with fresh media. The cells were cultured for an additional 48 hours followed by passaging to obtain a cell pellet. The pellet was resuspended in PBS and irradiated at 60 Gy for 20 minutes. Following irradiation, the cell solution was centrifuged, then resuspended in PBS and total cell number was counted using a hemocytometer. The cells were centrifuged and resuspended in freezing media (glioma media with 10% DMSO) at a concentration of 1 × 10^7^ cells/mL/cryovial. The cryovial was frozen in isopropanol chambers at - 80 C until use. On days 3, 7, and 14 following tumor-induction, cryovials were removed from −80 C and placed in 37 C water bath then resuspended in warm glioma media. Cells were centrifuged, washed in glioma media followed by another centrifugation. The resulting cell pellet was then resuspended in 1mL PBS with 20ng GM-CSF. Animals in the vaccination groups received a subcutaneous injection between the shoulder blades.

Following tumor induction, rats were monitored by lab member and University of Minnesota veterinary technicians three times daily, for 90-days post-tumor induction. Moistened lab chow was placed in a dish in the bedding to allow mice access to food and fluids. The point at which the animal was unable regain an upright posture, or demonstrate response to physical stimulation from the observer, or unable to lift their neck and head, the animal was declared moribund and immediately euthanized via Ketamine overdose (100mg/kg, *i.p.*) followed transcardial perfusion with ice-cold PBS and 4% paraformaldehyde.

### Data analysis and visualization

Survival data was analyzed using RStudio Environment (Version 1.0.153) using the Chi-Squared Kaplan-Meier method in the Survival package. Pairwise comparisons of each treatment group against untreated tumor-bearing mice in each study was corrected using Bonferroni correction for multiple comparisons. *In vitro* stimulation and IFN-γ detection statistical analysis was performed by one-way analysis of variance (ANOVA) with Tukey post hoc correction for multiple comparisons using GraphPad Prism 8 (GraphPad Software, San Diego, CA).

## Results

### GL261 cells are permissive to ZIKV infection

The ability for ZIKV to infect, replicate and release virus in GL261 cells was measured by plaque assay on Vero cells. An increase in virus particles within the media of infected cells was observed at one day post-infection (DPI) which remained elevated over time (Fig 1A). RNA isolated from lysed GL261 cells 3-DPI further confirmed the ability for ZIKV to infect GL261 cells as transcripts for the envelope (Fig 1B), non-structural 2 subunit (Fig 1C), and non-structure 5 subunit (Fig 1D) were observed in the infected samples. Finally, we assessed antiviral response gene expression following ZIKV infection in GL261 cells and a ZIKV non-permissive mouse microglia (BV2) (6). Cell lysates were collected at 1-DPI and 3-DPI for qRT-PCR analysis of transcripts related to endosomal (TLR3) and cytoplasmic (MDA5 and RIG-I) viral RNA detection (14). Relative to mock-infected cells, an increase in expression of TLR3 was observed at 1-DPI and 3-DPI, and increases in expression of MDA5 and RIG-I were observed at 3-DPI, which was not observed in BV2 cells (Fig 1E). These data suggest that, similar to previous studies, GL261 cells are permissive to ZIKV infection and propagation *in vitro*.

**Fig 1.**
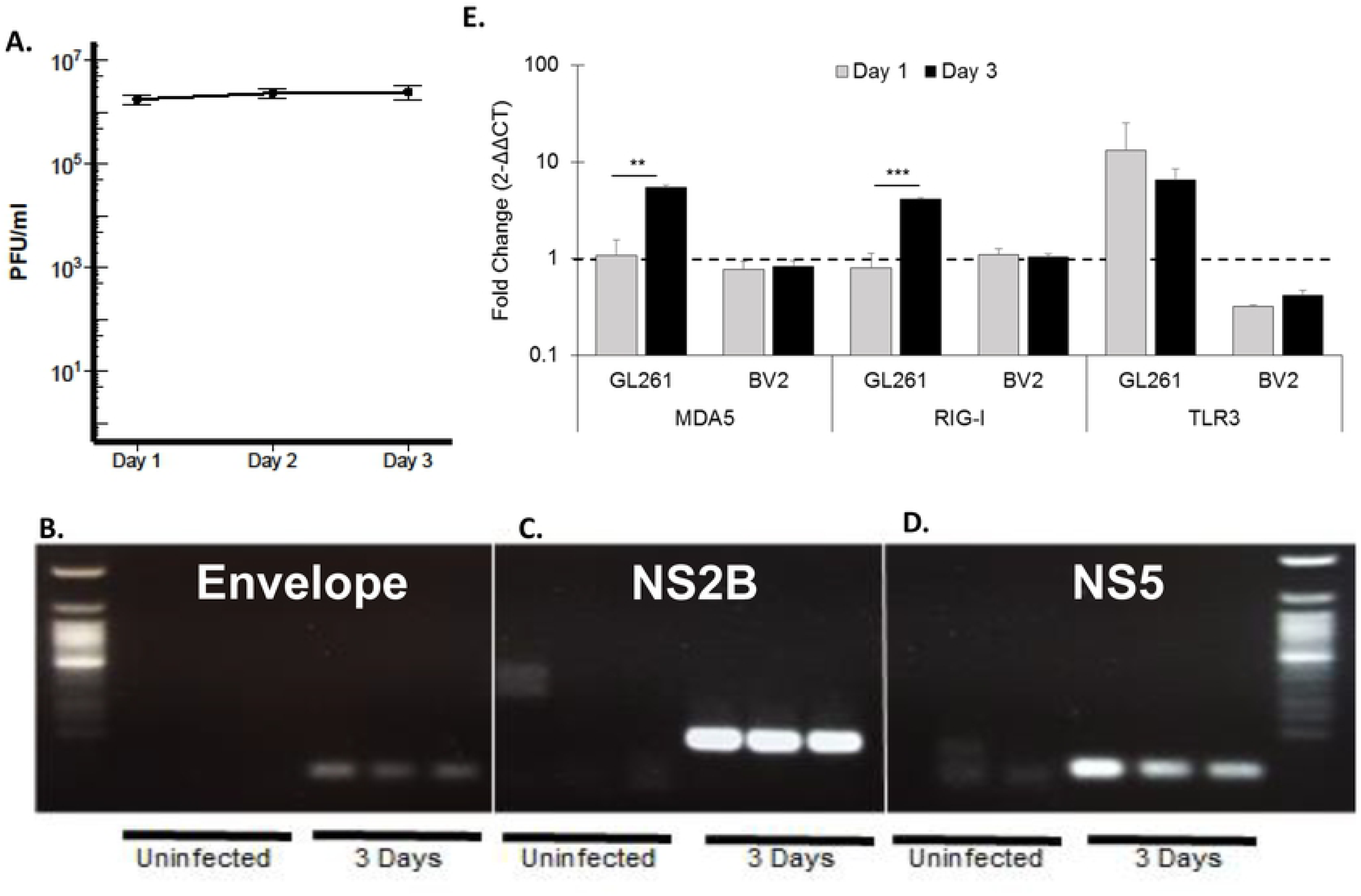
*In vitro* ZIKV infection of the murine GL261 glioma cell line. **A**) Plaque forming assay demonstrating permissiveness of ZIKV to infect and release virus, measured at 24-hour intervals following initial infection of 0.01 MOI (~2-3 × 10^4^ PFU). PCR and electrophoresis-based detection of ZIKV transcripts for Envelope (**B**), NS2B (**C**), and NS5 (**D**) 3-days following infection in GL261 cells. **E**) Expression of viral pattern recognition receptor (MDA5, RIG-I, and TLR3), in ZIKV infected cells at 1- and 3-days post-ZIKV infection, measured via qRT-PCR. Data represent mean ± SEM, n=3 each cell line/time point, (\**p* < 0.05; \*\**p* < 0.01; \*\*\**p* < 0.001).

### Oncolytic effects of ZIKV in experimental glioma

To assess the ability of ZIKV to improve OS, *in vivo*, immunocompetent C57BL/6J mice were implanted with a single-cell suspension of 3 × 10^4^ GL261 cells directly into the striatum followed immediately by *i.c.* injections of 5 × 10^4^, 5 × 10^6^, or 5 × 10^8^ PFU ZIKV at the same coordinates. Mice were monitored daily for a moribund state as a humane endpoint for overall survival (OS) up to 118 days post-tumor induction. No significant alteration in OS in any of the ZIKV treated GL261-mice was observed (Figs 2A-C). In untreated GL261-mice, a median survival of 38 days with an OS of 15.7% was observed (Table 1). Only in GL261-mice treated with the middle dose of ZIKV (5×10^6^ pfu) was a non-significant trend in median survival of 46 days and an OS of 33.3% observed.

**Table 1.** Descriptive statistics mouse tumor studies

| Group | n | Median Survival (days) | Overall Survival (%) | Pairwise *p* |
| --- | --- | --- | --- | --- |
| GL261 | 19 | 38 | 15.7 | --- |
| GL261 + $5 \times 10^4$ ZIKV | 18 | 39.5 | 11.1 | 0.85 |
| GL261 + $5 \times 10^6$ ZIKV | 9 | 46 | 33.3 | 0.21 |
| GL261 + 5x10<sup>8</sup> ZIKV | 9 | 36 | 0 | 0.29 |
| GL261 + 5x10<sup>4</sup> ZIKV + Vaccine | 14 | 42 | 35.7 | 0.32 |
| GL261 + GL261 Vaccine | 6 | 41 | 16.6 | 0.68 |
| SCID GL261 | 7 | 32 | 0 | --- |
| SCID GL261 + 5x10<sup>4</sup> ZIKV + Vaccine | 7 | 30 | 0 | 0.61 |

**Fig 2.**
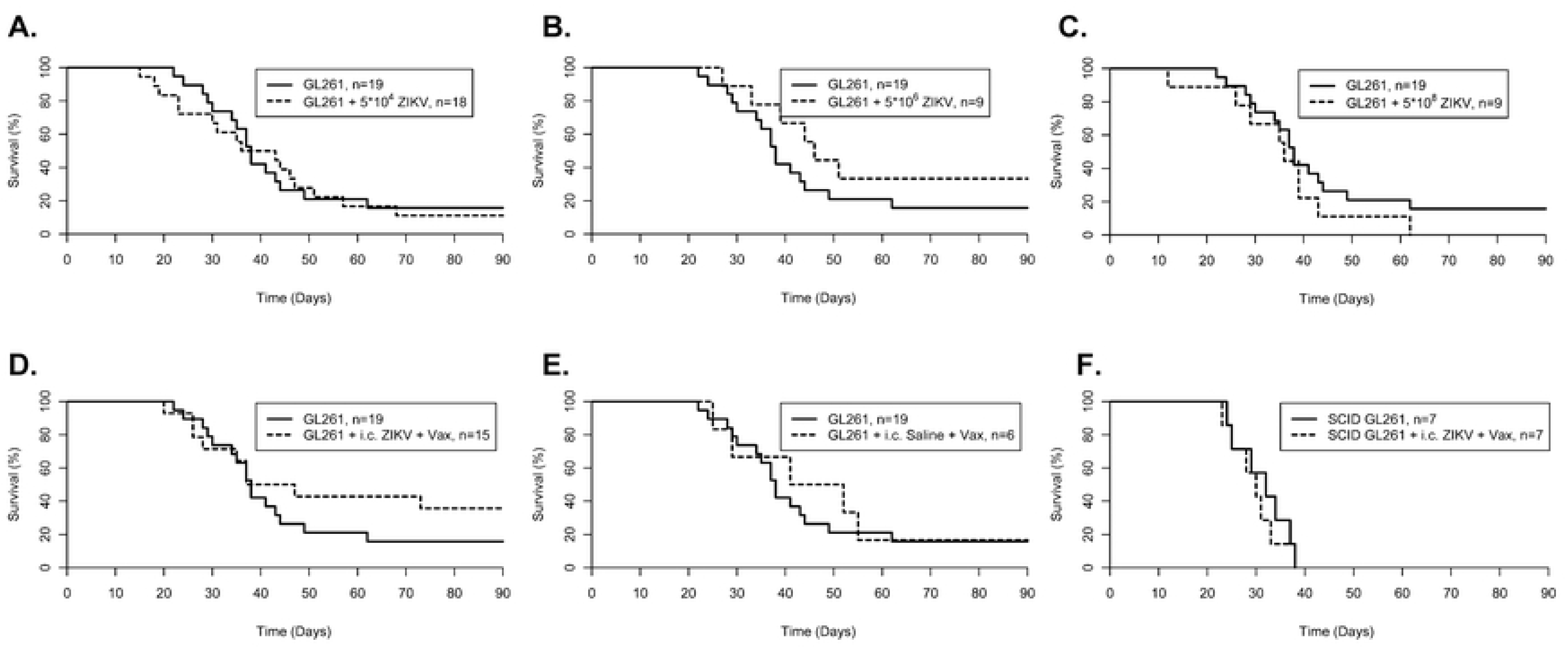
Survival plots in oncolytic study and first vaccine study. C57BL6/J mice implanted with murine GL261 glioma cells treated with intratumoral injection of ZIKV at 5×10^4^ (**A**), 5×10^6^ (**B**), 5×10^8^ (**C**) PFU, intratumoral injection of 5×10^4 PFU^ ZIKV coupled with peripheral vaccination of irradiated GL261 cells previously infected with ZIKV (GL261 ^+^ ZIKV ^+^ Vax; **D**), or peripheral vaccination of irradiated GL261 cells (GL261 ^+^ Vax; **E**) all relative to untreated GL261 implanted mice. **F**) NOD/SCID mice implanted with murine GL261 glioma cells treated with intratumoral injection of 5×10^4^ ZIKV coupled with peripheral vaccination of irradiated GL261 cells previously infected with ZIKV (GL261 ^+^ ZIKV ^+^ Vax) compared to untreated GL261 implanted NOD/SCID mice.

Similarly, we tested the oncolytic effect of ZIKV in a rat glioma model, where 5 × 10^4^ rat 9L cells were transplanted into in F344 rats and immediately treated with an intracranial injection of 5 × 10^4^ live ZIKV. Treatment of rats with *i.c.* injection of ZIKV alone did not improve OS (median survival of 32 days and OS of 0%) relative to untreated 9L tumor-bearing rats (median survival of 30 days and OS of 0%; Fig 3A; Table 2). These data indicate that ZIKV alone is not sufficient as a therapy for glioma.

**Table 2.** Descriptive statistics of rat tumor study

| Group | n | Median Survival (days) | Overall Survival (%) | Pairwise *p* |
| --- | --- | --- | --- | --- |
| 9L | 7 | 30 | 0 | --- |
| 9L + 5x10<sup>4</sup> ZIKV | 6 | 32 | 0 | 0.78 |
| 9L + 5x10<sup>4</sup> ZIKV + Vaccine | 6 | 29 | 16.6 | 0.38 |
| 9L + IR 5x10<sup>4</sup> ZIKV + Vaccine | 6 | n/a | 66.6 | 0.01* |
| 9L + 9L Vaccine | 6 | 50 | 33.3 | 0.02 |

**Fig 3.**
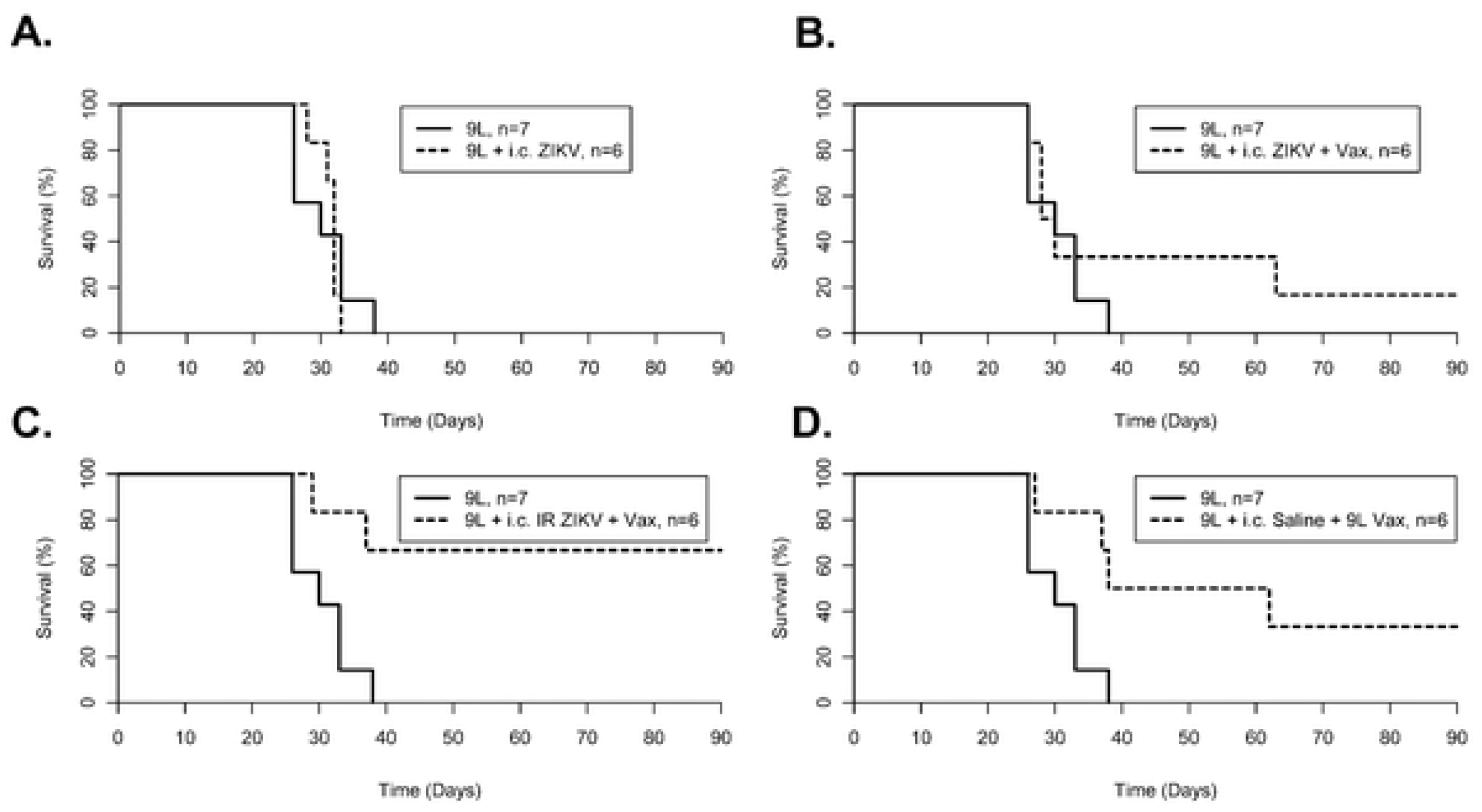
Survival plots in 9L rat model of glioma. F344 rats implanted with rat 9L glioma cells treated with either *i.c.* injection of 5×10^4^ live ZIKV (9L ^+^ ZIKV; **A**), intratumoral injection of 5×10^4^ live ZIKV coupled with peripheral vaccination of irradiated 9L cells previously infected with ZIKV (9L ^+^ ZIKV ^+^ Vax; **B**), intratumoral injection of 5×10^4^ irradiated ZIKV coupled with peripheral vaccination of irradiated 9L cells previously infected with ZIKV (9L ^+^ IR ZIKV ^+^ Vax; **C**), or peripheral vaccination of irradiated 9L cells (9L ^+^ 9L Vax; **D**) compared to untreated 9L implanted rats.

#### ZIKV & tumor vaccine co-therapy in experimental GLIOMA

Given that ZIKV was ineffective alone, we hypothesized that it could instead serve as an adjuvant to enhance the anti-tumor effects of a vaccine-based therapy. To test this, GL261-mice were treated with intracranial injections of ZIKV as well as subcutaneous vaccination (comprised of 5×10^6^ IR GL261 cells previously infected with ZIKV and 20ng of the dendritic cell adjuvant GM-CSF) on days 3, 7, and 14 post-tumor induction. OS of GL261-mice in the *i.c.* ZIKV plus vaccine-treated group was at 35.7% compared to 15.7% in the untreated controls and 16.6% in GL261-mice treated only with vaccines (Figs 2D-E; Table 1). Pairwise Kaplan-Meier analysis with Bonferroni correction, however, did not reach significance. In the 9L model of glioma, rats treated with live ZIKV coupled with a 9L-based vaccine had a non-significant increase in OS (Fig 3B; Table 2). Interestingly, a significant increase in OS was observed in 9L bearing rats treated with the combination of IR *i.c.* ZIKV and subcutaneous vaccines (OS of 66.6%; pairwise *χ*^2^ *=* 6.5, *p* = 0.011; Fig 3C; Table 2).

To evaluate the role of the immune system in this therapy, immunodeficient NOD-SCID mice were implanted with GL261 cells and subjected to the same ZIKV plus vaccine treatment. Relative to untreated NOD-SCID GL261-mice, no significant difference in OS was observed (Fig 2F; Table 1). The difference in response to treatment between the immunocompetent and immunodeficient mice supports the notion that the effects of *i.c.* ZIKV administration coupled with peripheral vaccination is mediated through an immune response.

To further investigate the role of the immune system in this adjuvant vaccine treatment paradigm, long-term survivors of the *i.c.* ZIKV plus vaccine-treated group were subjected to a second injection of GL261 cells approximately 114 days post-tumor-induction. Long-term survivors and age-matched naïve controls were implanted with either 3 × 10^4^ GL261 cells or received saline at the same coordinates. Seven days following rechallenge, brains were collected for flow cytometry (**S1 Fig**). Relative to saline-injected long-term survivors and age-matched naive tumor-implanted mice, we observed significant increases in the number of total T-cells (Fig 4A) and CD4^+^ T-cells (Fig 4B) in tumor-rechallenged long-term survivors. Memory CD4^+^ T-cells (Fig 4C) and CD69^+^ activated T-cells (Fig 4D), as well as CD11c^+^ dendritic cells and activated MHC II^+^ microglia also increased in number (Figs 4E-F). These results reflect a coordinated activation of immune cells in long-term survivors, suggesting that *i.c.* ZIKV and tumor vaccine immunotherapy enhances a tumor-specific T-cell response resulting in long-term immunity.

**Fig 4.**
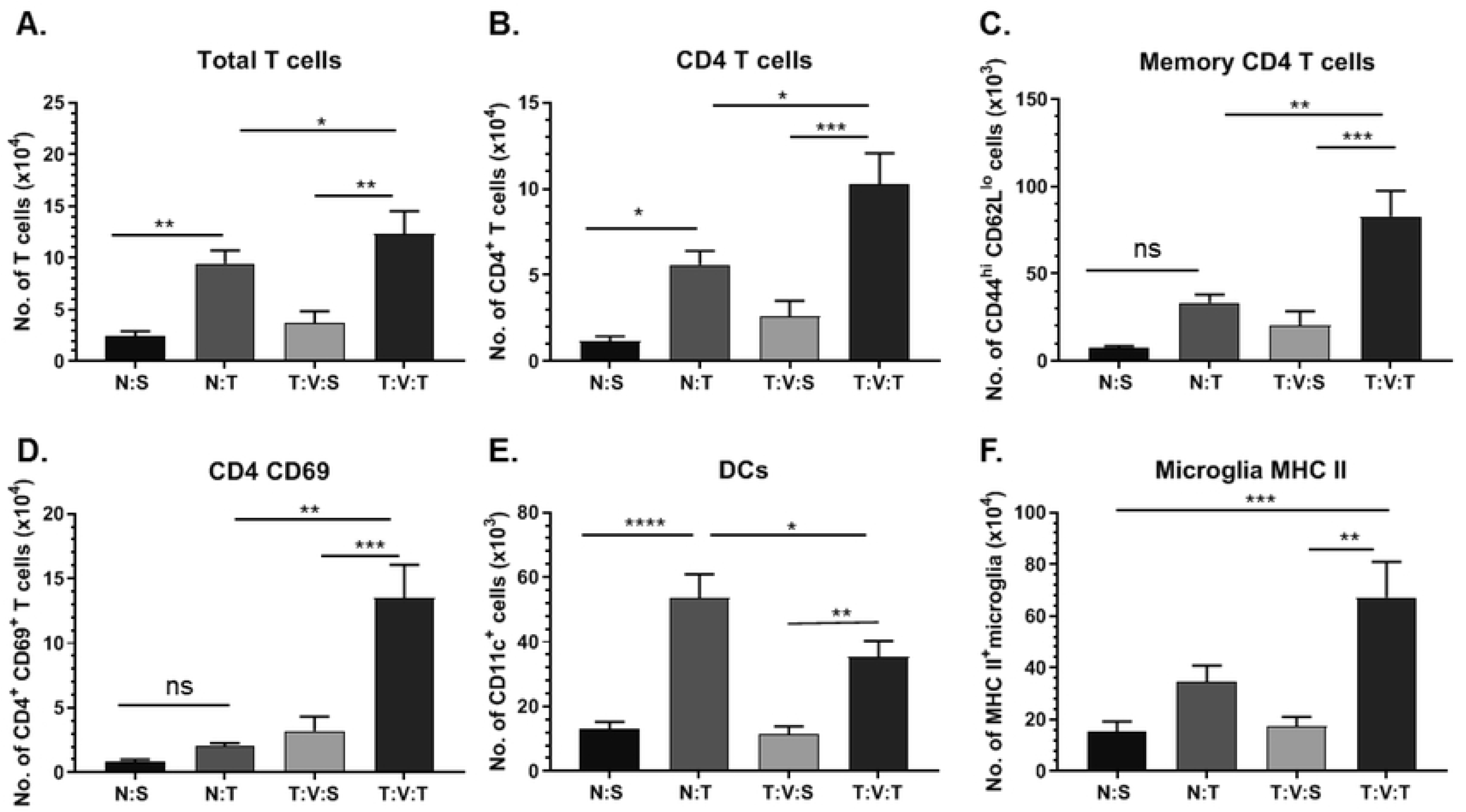
Phenotype of infiltrating and resident immune cells in tumor re-challenged mice. Flow cytometry analysis of mononuclear cells within the brains of long-term survivors of first vaccine treatment study 7 days following tumor re-challenge (T:V:T, *n*=5), long-term survivors 7 days following intracranial injection of saline (T:V:S, *n*=4), age-matched naïve-mice 7 days following intracranial tumor injection (N:T, *n*=8), or age-matched naïve-mice 7 days following intracranial saline injection (N:S, *n*=4). Total number of **(A)** total T-cells, **(B)** CD4^+^ T-cells, **(C)** CD4^+^ CD44^hi^ CD62L^lo^ memory T-cell, **(D)** activated CD4^+^ CD69^+^ T-cells **(E)** dendritic cells, and **(F)** activated MHC II^+^ microglia. Data represent mean ± SEM, (\**p* < 0.05; \*\**p* < 0.01; \*\*\**p* < 0.001).

### Irradiated ZIKV and tumor vaccine immunotherapy in a mouse glioma model

To increase the safety and efficacy of the protocol, we sought to match *i.c.* injections of our therapies to the peak T-cell response generated by peripheral vaccination. It has been reported that vaccinations in mice are expected to yield a peak T-cell proliferation approximately 10 days after the first vaccine injection (15), corresponding to day 13 in the first treatment paradigm. We, therefore, interrogated the effect of *i.c.* injections of our treatments on day 14. We also tested intracranial injection of IR GL261 cells previously infected with ZIKV based on results from the 9L rat model of glioma.

In this second paradigm, GL261-mice were administered subcutaneous vaccines on days 3, 7, and 21. In addition to subcutaneous vaccines, one group received *i.c.* injections of 5 × 10^4^ irZIKV immediately following tumor induction (day 0) and a second group on day 14. A third group received *i.c.* injection of IR GL261 cells previously infected ZIKV (*i.c.* vaccine) on day 14.

Similar to the first treatment paradigm, peripheral vaccinations of GL261-mice coupled with *i.c.* irZIKV immediately following tumor induction did not significantly improve median survival (35 days) or OS (14.3%), relative to untreated GL261-mice (Fig 5A; Table 3). Delaying *i.c.* irZIKV injection until day 14 significantly increased OS, relative to untreated mice (50%; pairwise *χ*^2^ = 11.1, *p* < 0.001; Fig 5B). Furthermore, treating GL261-mice with peripheral vaccinations coupled with *i.c.* vaccine on day 14 resulted in the greatest number of mice surviving long-term (75% OS; pairwise *χ*^2^ *=* 16.1, *p* < 0.001; Fig 5C).

**Table 3.** Descriptive statistics of second mouse tumor study

| Group | n | Median Survival (days) | Overall Survival (%) | Pairwise *p* |
| --- | --- | --- | --- | --- |
| GL261 | 8 | 34 | 0 | --- |
| GL261 + Vaccine + IR $5 \times 10^4$ ZIKV (D0) | 7 | 35 | 14.3 | 0.23 |
| GL261 + Vaccine + IR $5 \times 10^4$ ZIKV (D14) | 8 | 46 | 50.0 | <0.001* |
| GL261 + Vaccine + *i.c.* Vaccine (D14) | 8 | NA | 75.0 | <0.001* |

**Fig 5.**
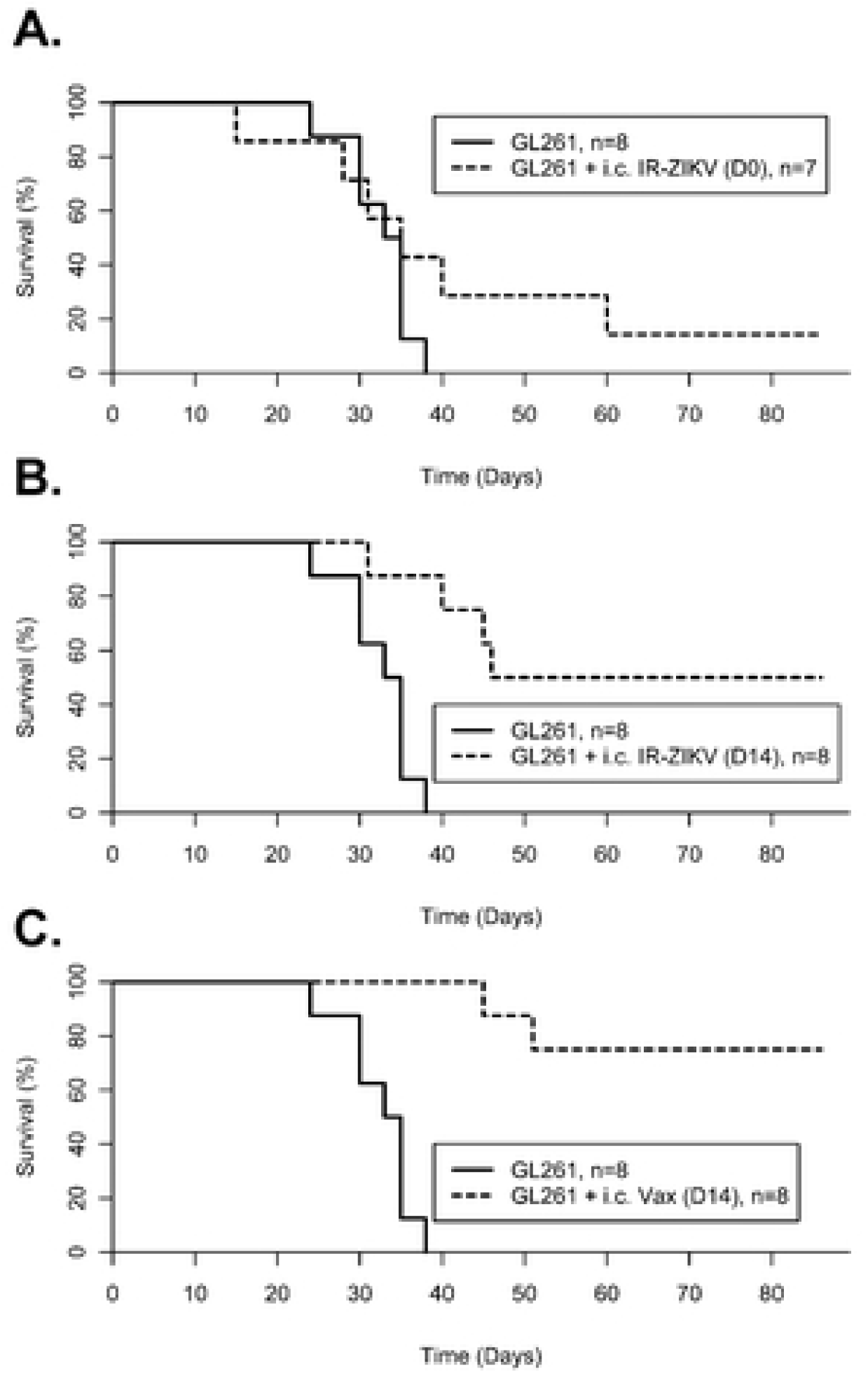
Survival plots of second mouse vaccine study. C57BL6/J mice implanted with murine GL261 glioma cells treated with intracranial injection of 5×10^4^ pfu irradiated ZIKV at Day 0 (**A**), intracranial injection of 5×10^4^ pfu irradiated ZIKV at Day 14 (**B**), or intracranial injection of 6×10^4^ irradiated GL261 cells previously infected with ZIKV (i.c. Vax) at Day 14 (**C**) all relative to untreated GL261 implanted mice.

To investigate the immune response of mice treated with peripheral vaccination coupled with *i.c.* vaccines, long-term survivors were rechallenged with GL261 cells approximately 86 days following tumor induction. The brains and cervical lymph nodes of animals were removed 7 days following rechallenge. Flow cytometry analysis of mononuclear cells isolated from brain tissue revealed treated rechallenged mice significantly increased levels of total T-cells within the brain (Fig 6A; **S2 Fig**), including CD4^+^ T-cells (Fig 6B) memory CD4^+^ T-cells (Fig 6C), and activated CD69^+^ CD4^+^ T-cells (Fig 6D), relative to naïve age-matched GL261-mice. Activated CD103^+^ resident memory T-cells within the brain were observed only in *i.c.* vaccinated mice re-challenged with tumor (Fig 6E). T-cells isolated from cervical lymph nodes were collected for functional assessment of antigen-specific IFN-γ production. Mice rechallenged with tumor following treatment with *i.c.* vaccination had increased levels of IFN-γ recall response when stimulated *ex vivo* with tumor lysate, but not when stimulated with ZIKV (Figure 5F). The results from this immunotherapy study further delineate the role of ZIKV as an adjuvant that can enhance tumor vaccines and is capable of improving OS to 75% in GL261-mice.

**Fig 6.**
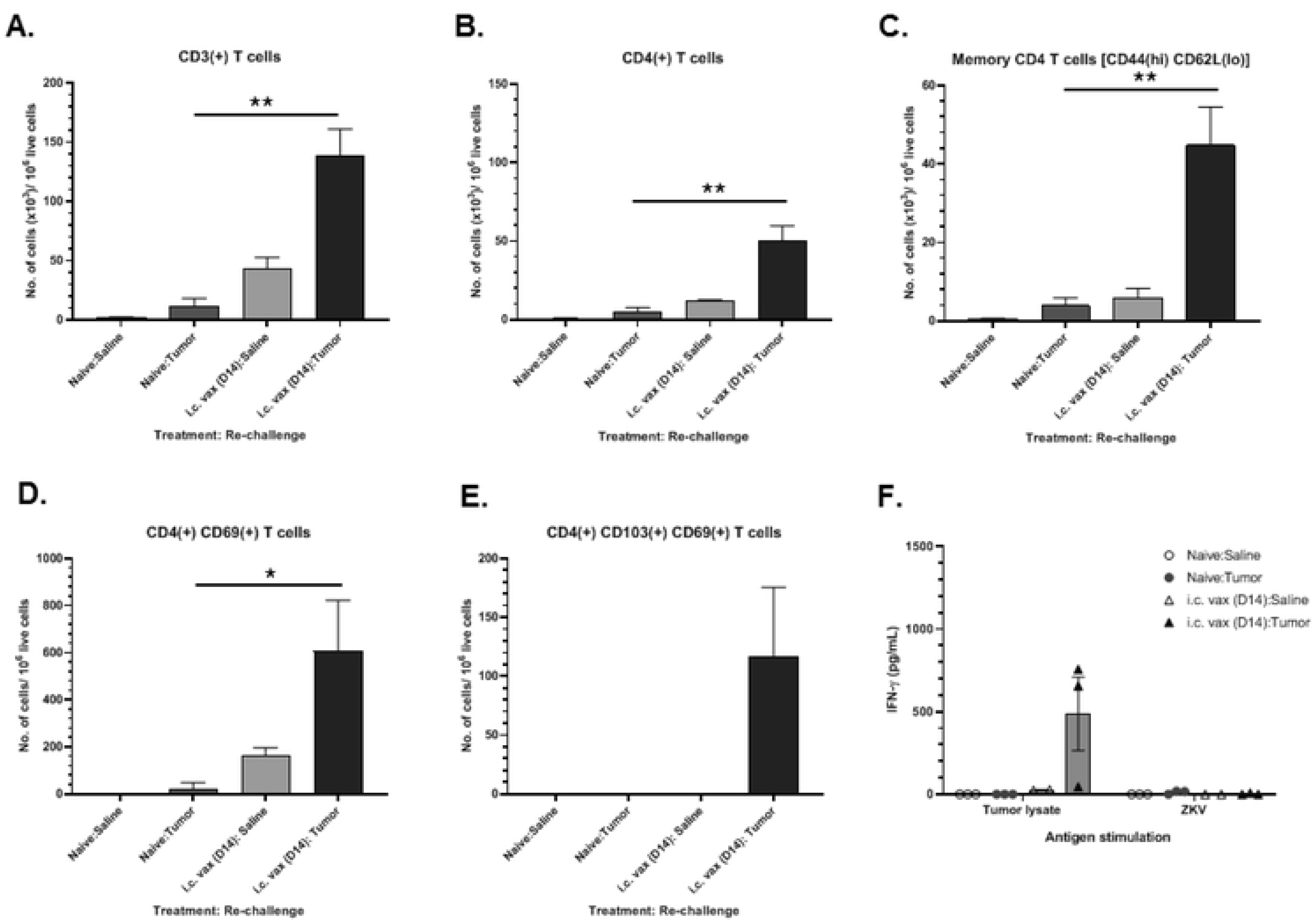
Phenotypes of infiltrating and resident immune cells and recall T-cell response in tumor re-challenged mice. Flow cytometry analysis of mononuclear cells within the brains of long-term survivors of second vaccine treatment study 7 days following tumor re-challenge (i.c. Vax: Tumor, *n*=3), long-term survivors 7 days following intracranial injection of saline (i.c. Vax: Saline, *n*=3), age-matched naïve-mice 7 days following intracranial tumor injection (Control: Tumor, *n*=3), or age-matched naïve-mice 7 days following intracranial saline injection (Control: Saline, *n*=3). Total number of **(A)** total CD3^+^ T-cells, **(B)** CD4^+^ T-cells, **(C)** CD4^+^ memory T-cell, **(D)** activated CD4^+^ T-cells, and **(E)** activated memory resident T-cells. **(F)** ELISA quantification of IFNγ secretion by lymphocytes isolated from cervical lymph nodes after 72 h *ex vivo* stimulation with tumor lysate or ZIKV in long-term survivors or age matched controls following re-challenge with tumor or saline. Data represent mean ± SEM, (\**p* < 0.05; \*\**p* < 0.01; \*\*\**p* < 0.001).

## Discussion

Virotherapy is an enticing approach for GBM as it exploits the propensity for viruses to infect and lyse tumor cells, leading to immune activation (16–19). In the present study, we observed that ZIKV alone had no therapeutic effects as an oncolytic virus in a mouse or rat glioma model, in contrast to work by other groups (6,8). It is likely that the strain of ZIKV and animal model used can account for the differences in oncolytic activity *in vivo*. Zhu and colleagues treated immunocompetent glioma-bearing mice with a Dakar strain of ZIKV which was serially passaged through a *Rag1*^−/-^ mouse, thereby increasing its virulence, specifically in mice (20), while Chen and colleagues treated immunodeficient tumor-bearing mice with a modified Cambodian strain (21).

While ZIKV was ineffective alone, combining virotherapy with other immunotherapy treatments is emerging as an effective preclinical paradigm for cancer treatment (22). Recently, Zhu and colleagues demonstrated that ZIKV infection of glioma stem cells *ex vivo* elicits an immune response including increased expression of genes related to adaptive immunity, antigen presentation, interferon response, and Toll-like receptor signaling pathways (23). Based on the potential for ZIKV to stimulate an immune response, we hypothesized that intratumoral ZIKV injections would penetrate the immunosuppressive tumor microenvironment and recruit T-cells generated by subcutaneous vaccinations to the tumor site to eliminate cancer cells. This was evident through the increase in OS in our second mouse study, where 50% of GL261-mice treated with *i.c.* irZIKV survived long-term. Furthermore, the intracranial administration of additional tumor antigen with ZIKV via *i.c.* vaccines produced an even greater OS, at 75%. We believe the enhanced OS in these two groups is due to the altered paradigm in which we targeted *i.c.* injections to peak T-cell response arising from subcutaneous vaccinations.

Successful outcomes following immunotherapies for the treatment of cancer have been linked to the generation of effector CD8^+^ and CD4^+^ T-cell responses, followed by the establishment of memory T-cell population capable of combatting re-emergence of the tumor (24). In the current study, we observed an increase in the total number of T-cells, CD4^+^ T-cells, and memory T-cells following tumor re-challenge in treated GL261-mice surviving long-term. CD4^+^ T-cells are crucial in orchestrating antitumor immunity, with growing evidence highlighting the role of CD4^+^ T-cells in combatting the immunomodulatory effects of tumors (25). In addition to the role of effector CD4^+^ T-cells, CD4^+^ memory T-cells are instrumental in protecting against future challenges and are the basis for successful vaccines (26). Our data, in addition to supporting the role CD4^+^ effector T-cells in driving an anti-glioma response, highlight the importance of memory CD4^+^ T-cells in providing defense against glioma reoccurrence. By eliciting a CD4^+^ T-cell response and generating memory T-cells selectively responsive to tumor antigen, ZIKV-based immunotherapy offers a promising avenue for providing long-term protection against glioma.

## Supporting information

**S1 Table.**
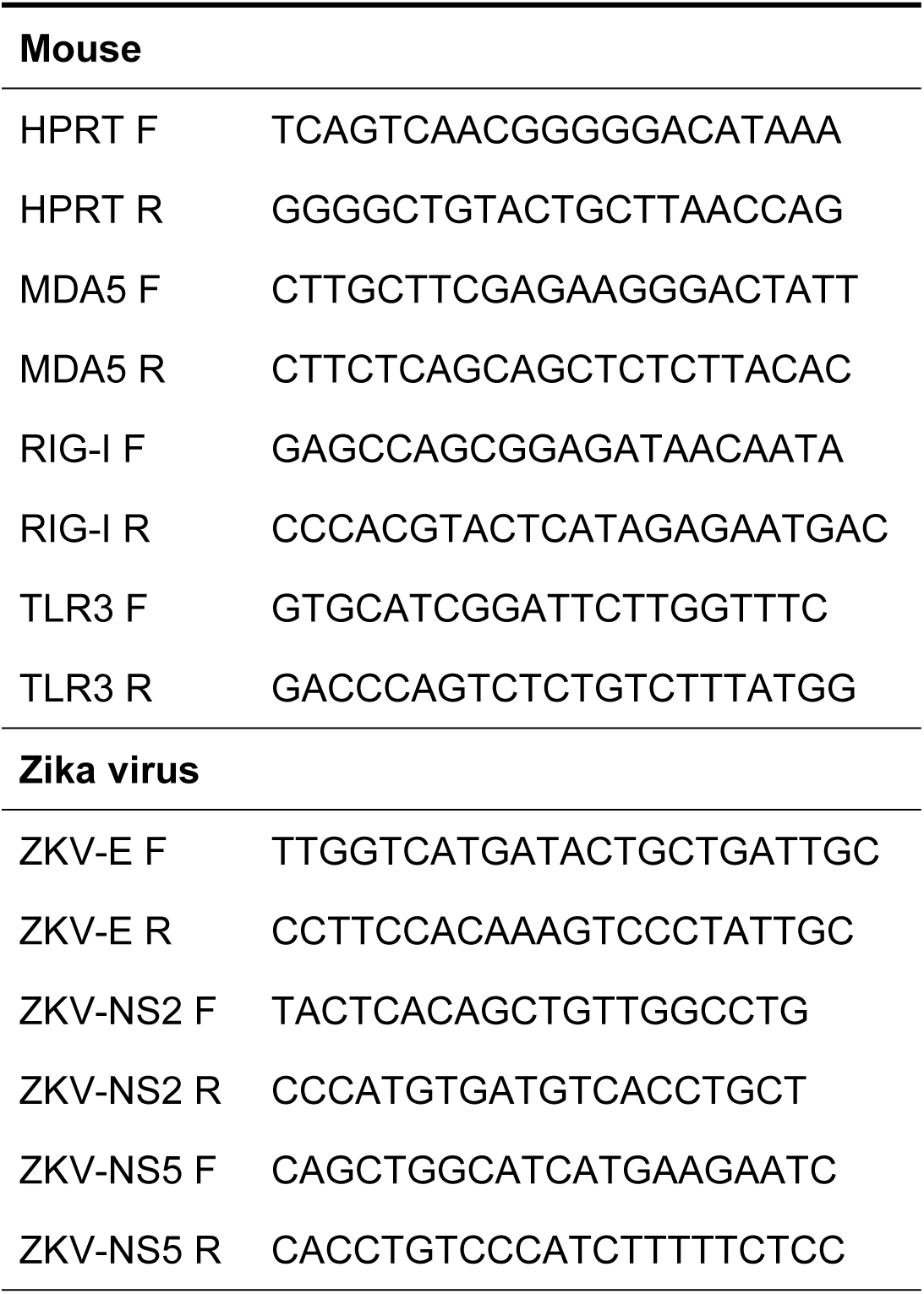
Primers for qRT-PCR Mouse

**S1 Fig. Flow cytometry analysis of immune cells in the brain following rechallenge of long-term survivors of first vaccine treatment study.** Mononuclear cells isolated from brain were analyzed by flowc ytometry as described in materials and methods. (A). Flow cytometry gating strategy to analyze immune cells in the brain. Debris (SSC-A vs FSC-A) and doublets (SSC-H vs SSC-A) were excluded and discrimination of live and dead cells were determined based on Live/Dead fixable Aqua dead cell staining. Live cells were gated to identify dendritic cells [DCs; CD45(hi) CD11c(^+^)], myeloid cells [CD45(hi) CD11b(^+^)], lymphoid cells [CD45(hi) CD11b(-)], and microglia [CD45(int) CD11b(^+^)]. CD45(hi) CD11b(-) cells were sub-gated to identify CD19(^+^) B cells and CD3(^+^) T-cells. Memory CD4 T-cells were identified as CD44(hi) CD62L(lo) cells and activated CD4 T-cells were identified based on CD69 expression. (B) Representative flow cytometry plots showing expression of memory and activation markers in different groups of mice. T:V:T-long-term survivors 7 days following intracranial injection of tumor; T:V:S-long-term survivors 7 days following intracranial injection of saline; N:T-age-matched naïve-mice 7 days following intracranial tumor injection; N:S-age-matched naïve-mice 7 days following intracranial saline injection.

**S2 Fig. Flow cytometry gating strategy to analyze immune cells in the brain following rechallenge of long-term survivors of second vaccine treatment study.** Mononuclear cells isolated from brain were analyzed by flow cytometry as described in materials and methods. Debris (SSC-A vs FSC-A) and doublets (SSC-H vs SSC-A) were excluded and live cells were gated to identify CD19(^+^) B cells and CD3(^+^) T-cells. Memory CD4 T-cells were identified as CD44(hi) CD62L(lo) cells, which are sub-gated based on CD103 expression as tissue resident memory T-cells.

**S1 File. Raw data.** Excel spreadsheet of raw data from mouse and rat survival studies, IFNγ secretion by lymphocytes, and CT values from qRT-PCR.

## References

1. Davis ME. Glioblastoma: Overview of Disease and Treatment. Clin J Oncol Nurs. 2016 Oct 1;20(5 Suppl):S2–8.

2. Bi WL, Beroukhim R. Beating the odds: extreme long-term survival with glioblastoma. Neuro Oncol. 2014 Sep;16(9):1159–60.

3. Russell SJ, Peng K-W, Bell JC. Oncolytic virotherapy. Nature Biotechnology. 2012 Jul 1;30(7):658–70.

4. Kaufman HL, Kohlhapp FJ, Zloza A. Oncolytic viruses: a new class of immunotherapy drugs. Nature Reviews Drug Discovery. 2015 Sep 1;14(9):642–62.

5. Geletneky K, Hajda J, Angelova AL, Leuchs B, Capper D, Bartsch AJ, et al. Oncolytic H-1 Parvovirus Shows Safety and Signs of Immunogenic Activity in a First Phase I/IIa Glioblastoma Trial. Molecular Therapy. 2017 Dec 6;25(12):2620–34.

6. Zhu Z, Gorman MJ, McKenzie LD, Chai JN, Hubert CG, Prager BC, et al. Zika virus has oncolytic activity against glioblastoma stem cells. J Exp Med. 2017 Oct 2;214(10):2843.

7. Zhu Z, Mesci P, Bernatchez JA, Gimple RC, Wang X, Schafer ST, et al. Zika Virus Targets Glioblastoma Stem Cells through a SOX2-Integrin αvβ5 Axis. Cell Stem Cell [Internet]. 2020 Jan 16 [cited 2020 Jan 27]; Available from: http://www.sciencedirect.com/science/article/pii/S1934590919304710

8. Chen Q, Wu J, Ye Q, Ma F, Zhu Q, Wu Y, et al. Treatment of Human Glioblastoma with a Live Attenuated Zika Virus Vaccine Candidate. MBio. 2018 Sep 18;9(5):e01683–18.

9. Baronti C, Piorkowski G, Charrel RN, Boubis L, Leparc-Goffart I, de Lamballerie X. Complete coding sequence of zika virus from a French polynesia outbreak in 2013. Genome Announc. 2014 Jun 5;2(3):e00500–14.

10. Bierle CJ, Fernández-Alarcón C, Hernandez-Alvarado N, Zabeli JC, Janus BC, Putri DS, et al. Assessing Zika virus replication and the development of Zika-specific antibodies after a mid-gestation viral challenge in guinea pigs. PLoS One. 2017 Nov 3;12(11):e0187720–e0187720.

11. Baer A, Kehn-Hall K. Viral Concentration Determination Through Plaque Assays: Using Traditional and Novel Overlay Systems. J Vis Exp [Internet]. 2014 Nov 4 [cited 2019 May 13];(93). Available from: https://www.ncbi.nlm.nih.gov/pmc/articles/PMC4255882/

12. Pino PA, Cardona AE. Isolation of brain and spinal cord mononuclear cells using percoll gradients. J Vis Exp. 2011 Feb 2;(48):2348.

13. Hamel R, Dejarnac O, Wichit S, Ekchariyawat P, Neyret A, Luplertlop N, et al. Biology of Zika Virus Infection in Human Skin Cells. J Virol. 2015 Sep;89(17):8880–96.

14. Pichlmair A, Reis e Sousa C. Innate Recognition of Viruses. Immunity. 2007 Sep 21;27(3):370–83.

15. Badovinac VP, Messingham KAN, Jabbari A, Haring JS, Harty JT. Accelerated CD8+ T-cell memory and prime-boost response after dendritic-cell vaccination. Nature Medicine. 2005 Jul 1;11(7):748–56.

16. Gromeier M, Lachmann S, Rosenfeld MR, Gutin PH, Wimmer E. Intergeneric poliovirus recombinants for the treatment of malignant glioma. Proc Natl Acad Sci U S A. 2000 Jun 6;97(12):6803–8.

17. Jiang H, Clise-Dwyer K, Ruisaard KE, Fan X, Tian W, Gumin J, et al. Delta-24-RGD oncolytic adenovirus elicits anti-glioma immunity in an immunocompetent mouse model. PLoS One. 2014 May 14;9(5):e97407–e97407.

18. Foreman PM, Friedman GK, Cassady KA, Markert JM. Oncolytic Virotherapy for the Treatment of Malignant Glioma. Neurotherapeutics. 2017 Apr;14(2):333–44.

19. Kaufman HL, Kohlhapp FJ, Zloza A. Oncolytic viruses: a new class of immunotherapy drugs. Nature Reviews Drug Discovery. 2015 Sep 1;14(9):642–62.

20. Gorman MJ, Caine EA, Zaitsev K, Begley MC, Weger-Lucarelli J, Uccellini MB, et al. An Immunocompetent Mouse Model of Zika Virus Infection. Cell Host & Microbe. 2018 May 9;23(5):672–685.e6.

21. Shan C, Xie X, Muruato AE, Rossi SL, Roundy CM, Azar SR, et al. An Infectious cDNA Clone of Zika Virus to Study Viral Virulence, Mosquito Transmission, and Antiviral Inhibitors. Cell Host & Microbe. 2016 Jun 8;19(6):891–900.

22. Suzuki M. Partners in Crime: Combining Oncolytic Viroimmunotherapy with Other Therapies. Mol Ther. 2017 Apr 5;25(4):836–8.

23. Zhu Z, Mesci P, Bernatchez JA, Gimple RC, Wang X, Schafer ST, et al. Zika Virus Targets Glioblastoma Stem Cells through a SOX2-Integrin αvβ5 Axis. Cell Stem Cell. 2020 Feb 6;26(2):187–204.e10.

24. Amsen D, van Gisbergen KPJM, Hombrink P, van Lier RAW. Tissue-resident memory T cells at the center of immunity to solid tumors. Nature Immunology. 2018 Jun 1;19(6):538–46.

25. Lai Y-P, Jeng C-J, Chen S-C. The Roles of CD4+ T Cells in Tumor Immunity. ISRN Immunology. 2011;2011:6.

26. MacLeod MKL, Clambey ET, Kappler JW, Marrack P. CD4 memory T cells: what are they and what can they do? Semin Immunol. 2009 Apr;21(2):53–61.

